## Supplemental Information for "Mostly-monocular responses and other visual functions in a multiscale network model of Macaque V1"

June 17, 2026

#### S1 Network structure and model parameters for the CG model

Here we describe in detail the CG model of Layer 4C$\alpha$ presented in the main text. The model is guided by the mechanistic network models in [1–3] and is an improved and expanded version of the CG model in [4]. We begin by describing the structure of the basic CG model, which was used to produce Figs. 2–3, then address its extension to binocular input (Figs. 4–7). The main difference is the inclusion of ocular dominance columns (ODCs): in the basic CG model, all visual input comes from one eye, and the L4 network architecture only depends on the distance between cells. In contrast, in the two-eye model, visual input originates from two eyes, and the L4 network architecture is modulated by ODCs.

##### S1A Model description

Neurons in L4 receive direct input from three external sources: feedforward input from LGN, feedback from layer 6 (abbrev. L6), and an "ambient" source that represents all top-down inputs not otherwise modeled (e.g., neuromdulatory inputs); see Fig. 1A in the main text.

###### Preliminary model without ODCs

**L4, the principal layer.** We begin by describing a biologically detailed network model of L4, followed by the coarse-graining procedure.

We divide L4 into *hypercolumns* (HCs) consisting of cells that share the same receptive fields. Each HC occupies a square area of $0.5 \times 0.5\,\text{mm}^2$, containing 3200 excitatory (E) cells and 800 inhibitory (I) cells uniformly distributed. L4 can also be subdivided into *orientation domains*, collections of cells that respond preferentially to similar orientations. Our model consists of $4 \times 4$ HCs depicted as squares in Fig. 1A; *intended orientation preferences* in different domains are marked with parallel bars. Connections between neurons within L4 do not respect HCs or orientation domains; the actual receptive field and preferred orientation of a L4 neuron are consequences of its thalamocortical input (discussed in Sect. S2) and interaction with other cortical neurons.

We distinguish between two subpopulations of E-cells in L4, *simple* (S) (comprising $\sim 70\%$ of E-cells) and *complex* (C) [5, 6]. These two kinds of cells receive different thalamocortical input (see below). As complex cells have significantly higher firing rates than simple cells and fewer LGN inputs, we have modified the connectivity within L4 to give them a larger set of presynaptic E-cells.

*Network design.* The structure and connectivity of this layer are as in [1–3]. Briefly, about 80% of the neurons are Excitatory (E); the rest are Inhibitory (I) and assumed to be basket cells [7]. We assume E and I-neurons are located on two square lattices on a 2D surface. Anatomical facts of the following type are incorporated: we model the set of presynaptic type $R$-cells to a type $Q$ cell ($Q, R \in \{E, I\}$) as a truncated Gaussian

$$P^{QR}(r) = \begin{cases} p^{QR}_{\text{peak}} \cdot \exp\left(-\frac{r^2}{\sigma_R^2}\right), & r < r^{\text{L4}} \\ 0, & r \geqslant r^{\text{L4}} \end{cases}$$

Here, $\sigma_R$ and $r_{\text{L4}}$ represent the spatial scale and truncation radius, respectively. As E-cells have longer axonal reach, we set $\sigma_R = 0.2/\sqrt{2}\,\text{mm}$ for excitatory (E) cells and $0.125/\sqrt{2}\,\text{mm}$ for inhibitory (I) cells. For both cell types, $r_{\text{L4}} = 0.36\,\text{mm}$. The anatomical data comes from [8, 9]. Peak connection probability among E-cells is $p^{EE}_{\text{peak}} \sim 15\%$, while peak connection probabilities for E→I, I→E, and I→I are much higher, at about 60% [10, 11]. Periodic boundaries are employed in both coarse-grained (CG) models to accommodate projections extending beyond the patch's boundaries. This ensures that cells located near the boundaries receive inputs from the same number of presynaptic neurons as cells situated at the center of the patch.

E-to-E connection probabilities are further modified to reflect the distinction between S and C-cells. Experimental data reveals that C-cells receive much more recurrent input and have higher firing rates than S-cells. To reproduce these features, previous studies assigned larger numbers of presynaptic E-cells (from both L4 and L6) and fewer LGN inputs to C-cells. Conversely, S-cells are assigned fewer presynaptic E-cells and more LGN inputs [1]. On average, S-cells receive inputs from 133 S, 57 C, and 85 I cells within L4; C-cells receive 179 S, 76 C, and 85 I inputs; and I-cells receive 591 S, 253 C, and 85 I inputs. These values are rounded averages derived from the anatomical assumptions above.

*Coarse-graining.* Following [4], we divide each HC into $10 \times 10$ "pixels," or local populations, with distinct inputs and outputs (see Fig. 1B). Each pixel contains a local circuit of $\sim 32$ E-cells and $\sim 8$ I-cells. Modeled are 3 cell types (simple-E, complex-E, and I) and 3 sources of "external" inputs (LGN, L6, and "ambient").

Interaction kernels among pixels are as defined by aggregating the average connectivity among neural populations, considering contributions from itself and nearby pixels. Specifically, for pixels $p$ and $p'$, the average number of type-$R$ cells in pixel $p'$ presynaptic to a type-$Q$ cell in pixel $p$ is represented by the L4 synaptic interaction kernel $C_{Q\leftarrow R;p\leftarrow p'}$ ($Q, R \in \{S, C, I\}$). The total number of L4 type-$R$ cells presynaptic to a type-$Q$ cell is

$$N^{QR} = \sum_{p' \in \mathscr{P}} C_{Q\leftarrow R;p\leftarrow p'}, \tag{1}$$

where $\mathscr{P}$ is the set of all pixels in the CG model. For more details of $C_{Q\leftarrow R;p\leftarrow p'}$, we refer readers to Eqs. (2) and (3) and Fig. 1 in the Supporting Information of [4].

Below we discuss in some detail three novel aspects of the present model that are significant improvements from the CG model in [4]. They are the feedforward and feedback actions of LGN and L6, and the depression of I-cells.

**Feedforward input from LGN.** Compared to [4], the modeling of thalamocortical input in this paper is both biologically more accurate and more flexible, able to respond to a larger collection of visual stimuli, e.g., gratings at different contrasts. Here we briefly highlight the differences and defer details to Sect. S2.

We model the dynamics of individual LGN cells as was done in [1, 3]. Briefly, there are two kinds of LGN cells, ON and OFF, responding to increments and decrements in luminance. Spontaneous LGN firing rates are $\sim 20$ spikes/s, rising with contrast to a mean firing rate of $45-50$ spikes/s or a peak firing rate of $> 100$ spikes/s when driven by a 10 Hz drifting grating at high contrast. The wiring between L4 cells and LGN follows [3].

The key difference between this paper and [4] lies in the *firing patterns* of the *pooled LGN input to a simple V1 cell*. This matters because single LGN cells are not orientation specific, so that the number of LGN spikes received by a V1 cell is independent of the orientation of the grating, and as a result it is not enough to simply attribute a mean firing rate to LIF neurons in the CG model. To accurately model responses of downstream L4 cells to gratings of different orientations, we need to convey information on firing patterns. Whereas square waves were used in [4] for simplicity, here we use actual spike trains to elicit more realistic responses. This is done by simulating in advance the responses of LGN to drifting gratings and recording their pooled spike trains to typical V1 cells (details in Sect. S2).

**Feedback from L6.** In the main text, we argued that L6-to-L4 feedback is simply a reflection of local L4 dynamics, describable as a local weighted average of L4 firing rates, with weights chosen to fit known properties of L6. This is implemented in the CG model as follows: given a firing rate configuration $f$ (see Materials & Methods), we introduce for each pixel $p$ a number $r(p)$ representing the local E-firing rate at $p$, defined to be a weighted average of the L4 E-firing rates (weighted average of S and C rates) in a neighborhood of $\sim 3 \times 3$ pixels centered at $p$. The kernel $K$ for this weighted average is derived from a Gaussian with standard deviation $\sigma = 37.5\,\mu$m (0.75 pixels) truncated at a radius of 1.5 pixels; see [12] and also [2]. This yields the $3 \times 3$ weight matrix

$$k = \begin{bmatrix} 0.058 & 0.117 & 0.058 \\ 0.117 & 0.300 & 0.117 \\ 0.058 & 0.117 & 0.058 \end{bmatrix} \tag{2}$$

In addition, to model synaptic depression at high firing rates, we introduce a *L6 response function* $R$, with the property that each L6 cell at pixel $p$ that projects to L4 is assumed to fire $R(r)$ spikes/sec when the local firing rate at pixel $p$ is equal to $r$. The graph of $R$ used is shown in Fig. S1; it is monontonically increasing, mostly linear with slope $\approx 2$, following data on L4 and L6 firing rates (see, e.g., [2] and references therein). The total number of L6 spikes received by a cell in pixel $p$ is then given by $m^{QL6} \times R(r(p))$, where $m^{QL6}$ is the number of presynaptic L6 cells for L4 cells of type $Q$, $Q \in \{S, C, I\}$ (see specific numbers below).

Before moving on, we point out that, on average, each S-cell in the CG model receives inputs from 4.5 LGN cells and 45 L6E cells; each C-cell receives inputs from 1.5 LGN and 55 L6E cells; each I-cell receives inputs from 4 LGN and 150 L6E cells.

**Depression of I-cells.** A novel feature of our network not in [4] is the depression of I-cells. It is a well established fact that I-cells in L4 are predominantly of PV type [13], and I-cells of PV-type depress when their firing rates are high, meaning their spikes become less potent. In our CG model, this translates into a decrease of the number of I-spikes by certain percentages. This is done through the creation of an *I-depression factor* $s(\cdot)$, where $s(x) \cdot x$ is the effective number of I-spikes when I-firing rate is $x$ sp/s. That

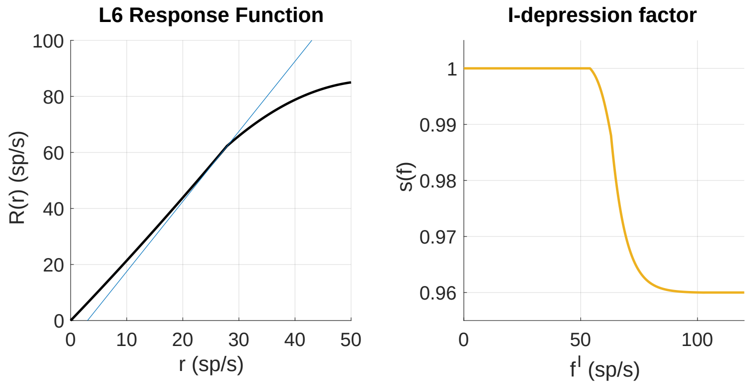


Figure S1: L4 and L6 response curves. Left: L6 response curve $R(r)$ (thick black line), and a reference curve with slope 2.5 (thin blue line). Right: L4 I-depression factor $s(\cdot)$.

is, when updating firing rate configuration $f$ (see Sect. S3B), if $x$ is the I-firing rate at pixel $p$, we simply replace it by $s(x)\cdot x$. A graph of $s(\cdot)$ is shown in the right panel of Fig. S1; note $s(x) = 1$ for $x < 54$.

##### Extension to two-eye model

The two-eye model differs from the basic CG model in the following ways. First, the two-eye model features an expanded cortical patch, comprising $8 \times 4$ HCs, with each row representing an ODC. Second, the LGN input to each ODC comes from a single eye and not from the opposite eye, a fact that has consequences for how we compute pixel responses (see Sect. S3). Lastly, and most critically, we adjust the L4 synaptic interaction kernels $C_{Q\leftarrow R;p\leftarrow p'}$ based on the values of $\mathbf{P}(Crs)$, $\mathbf{P}(Rfl)$, and $\mathbf{P}(Abs)$ (with $\mathbf{P}(Crs)+\mathbf{P}(Rfl)+\mathbf{P}(Abs) = 1$): when the projection $p \leftarrow p'$ crosses an ODC boundary, we set

$$\hat{C}_{Q\leftarrow R;p\leftarrow p'} = C_{Q\leftarrow R;p\leftarrow p'} \cdot \mathbf{P}(Crs), \tag{3a}$$

$$\hat{C}_{Q\leftarrow R;p\leftarrow p''} = C_{Q\leftarrow R;p\leftarrow p''} + C_{Q\leftarrow R;p\leftarrow p'} \cdot \mathbf{P}(Rfl), \tag{3b}$$

where the pixel $p''$ is a reflection of pixel $p'$ about the ODC boundary separating $p$ and $p'$, and use $\hat{C}_{Q\leftarrow R;p\leftarrow p'}$ in place of $C_{Q\leftarrow R;p\leftarrow p'}$ in the model; see, e.g., Eq. (1) above.

All other details are the same for the basic and two-eye models. In particular, L6 projections are insensitive to ODC boundaries.

#### S1B Responses of pixels

Within each pixel, cells of the same type are assumed to receive the same input and thus yield the same as, both measured by firing rates. We associate to each pixel $p$ a tuple of firing rates, $(f_p^S, f_p^C, f_p^I)$, one for each cell type. We compute firing rates using the leaky integrate-and-fire model:

$$\frac{dV}{dt} = -\frac{1}{\tau_L^Q} V - g_E^Q(t)(V - V^E) - g_I^Q(t)(V - V^I)\,, \tag{4}$$

where $V$ is the membrane voltage (nondimensionalized to $[0,1]$), the conductances satisfy

$$
\begin{aligned}
g_E^Q(t) \;=\;& \underbrace{S^{\text{QLGN}}\sum_{k=1}^{\infty} G_{\text{ampa}}(t-t_k^{\text{lgn}})}_{\text{(I) LGN}} + \underbrace{S^{\text{Qamb}}\sum_{k=1}^{\infty} G_{\text{ampa}}(t-t_k^{\text{amb}})}_{\text{(II) ambient}} \\
& \underbrace{+S^{\text{QL6}}\sum_{k=1}^{\infty}\left[\rho_{\text{ampa}}^{Q} G_{\text{ampa}}(t-t_k^{\text{L6}}) + \rho_{\text{nmda}}^{Q} G_{\text{nmda}}(t-t_k^{\text{L6}})\right]}_{\text{(III) L6}} \qquad (5) \\
& \underbrace{+S^{QE}\sum_{k=1}^{\infty}\left[\rho_{\text{ampa}}^{Q} G_{\text{ampa}}(t-t_k^{\text{L4}E}) + \rho_{\text{nmda}}^{Q} G_{\text{nmda}}(t-t_k^{\text{L4}E})\right]}_{\text{(IV) L4E}} .
\end{aligned}
$$

and

$$
g_I^Q(t) \;=\; S^{QI}\sum_{i=1}^{\infty} G_{\text{gaba}}(t-t_i^{\text{L4}I}). \qquad (6)
$$

Terms I-IV correspond to sources of excitation from LGN, ambient, L6, and L4E, respectively. The associated synaptic weights are denoted by $S^{\text{Qsource}}$ (precise values are given in Sect. S1C), while $t_k^{\text{source}}$ denote spike arrival times. Spikes from (recurrent) E-to-E connections within L4 are subject to synaptic failure: each such spike is dropped with 20% probability, independently of each other. The factors $\rho_{\text{ampa,nmda}}^{Q}$ are the proportions of AMPA and NMDA receptors activated by each E-spike. The model contains two types of excitatory currents, AMPA and NMDA, which elevate $g_E^Q(t)$ for different amounts of time: $G_{\text{ampa}}(\cdot)$ persists for a few milliseconds, whereas $G_{\text{nmda}}(\cdot)$ extends roughly 80 ms.

Details of LGN input spike trains are given in Sect. S2. The ambient input is assumed to be independent Poisson processes; rates are given in Sect. S1C. L4 and L6 inputs, i.e., $\{t^{\text{L4E}}\}$, $\{t^{\text{L4I}}\}$, and $\{t^{\text{L6}}\}$, are modeled by Poisson processes; their rates must be determined iteratively using the CG model, as detailed in Sect. S3B.

#### S1C Parameters for the CG Model

Here are the synaptic weights and neuronal parameters for the CG model [4]:

**Synaptic Coupling Weights**

- Excitatory-to-excitatory: $S^{EE} = 0.024$
- Inhibitory-to-inhibitory: $S^{II} = 0.120$
- Inhibition-to-excitation ratio: $S^{EI}/S^{EE} = 1.88$
- Excitation-to-inhibition ratio: $S^{IE}/S^{II} = 0.131$
- Ambient inputs: $S^{E\text{amb}} = S^{I\text{amb}} = 0.01$, $F^{S\text{amb}} = 600$ Hz, $F^{C\text{amb}} = 750$ Hz, $F^{I\text{amb}} = 435$ Hz.

**LGN and L6 projections to L4**

- LGN coupling weights: $S^{\text{ELGN}} = 2\cdot S^{EE}$, $S^{\text{ILGN}} = 2\cdot S^{\text{ELGN}}$
- L6 coupling weights: $S^{E\text{L6}} = \frac{1}{3}\cdot S^{EE}$, $S^{I\text{L6}} = \frac{1}{3}\cdot S^{IE}$

– Numbers of presynaptic L6 cells: $m^{SL6} = 45$, $m^{CL6} = 55, m^{IL6} = 150$

**Neuronal Physiology**

– Reset potential: $V_{\text{rest}} = 0$, spiking threshold: $V_{\text{th}} = 1$

– Reversal potentials: $V_E = \frac{14}{3}$, $V_I = -\frac{2}{3}$

– Leakage timescales: $\tau_L^E = 20\,\text{ms}$, $\tau_L^I = 16.7\,\text{ms}$

– Synaptic failure probability: $p_{\text{fail}} = 0.2$

– Refractory period: $\tau_{\text{ref}} = 2\,\text{ms}$

**Postsynaptic conductances:**

$$G_s(t) = \frac{1}{\tau_s^{\text{decay}} - \tau_s^{\text{rise}}} \left( e^{-t/\tau_s^{\text{rise}}} - e^{-t/\tau_s^{\text{decay}}} \right),$$

where $(\tau_s^{\text{rise}}, \tau_s^{\text{decay}})$ stand for the time scales of activation/inactivation of synapse type $s =$ ampa, nmda, gaba; the time constants used here are

– AMPA: $(\tau_{\text{rise}}, \tau_{\text{decay}}) = (0.5, 3)\,\text{ms}$

– NMDA: $(\tau_{\text{rise}}, \tau_{\text{decay}}) = (2, 80)\,\text{ms}$

– GABA: $(\tau_{\text{rise}}, \tau_{\text{decay}}) = (0.5, 5)\,\text{ms}$

#### S2 Simulation of LGN input to L4 cells

Here we describe in detail how we simulate LGN inputs for the purpose of precomputing L4 cell responses. As argued in the main text and in Sect. S1A, to faithfully model orientation selectivity and other basic V1 functions, we need to reproduce not just LGN inputs with correct mean firing rates but also their *temporal spike patterns*. As an example, consider a simple V1 cell in the vertical-preferring domain receiving input from 4 LGN cells. Following [1, 3], these 4 LGN cells typically have the following spatial configuration: they are aligned in two vertical columns of ON and OFF, with prescribed spacings between ON and OFF, as shown in Fig. S2. The top panel of Fig. S3 shows the response of a single LGN cell to a drifting grating with vertical bars (i.e., 0°) with spatial frequency (SF) = 2.5 cycles/deg (c/d) and temporal frequency (TF) = 10 Hz, at maximum contrast. The bottom panel shows the responses (in spikes/sec) of the 4 LGN cell-configuration pooled together. The pooled LGN spike trains would be much more temporally homogeneous when driven by orghogonal gratings (see, e.g., the bottom panel of Fig. 1D in the main text), though *time-averaged firing rates are essentially the same for the two gratings*.

In this paper, we are primarily interested in steady state responses of L4 local populations when the eye is presented with drifting gratings that are large enough to cover the visual field for the entire part of V1 being modeled. Under these conditions, only configurations of LGN afferents to L4 cells (and not their specific locations in LGN space) matter. It is sufficient to compute pooled spike trains like those in the example above for (i) all relevant LGN configurations, and (ii) visual stimuli described by a 4-dimensional space consisting of the *orientation*, *contrast*, *SF*, and *TF* of drifting gratings.

Note that the mean response of an $\alpha$-preferring local population to a grating with orientation $\theta$ can be equated with that of a vertical-preferring OD to a $(\theta - \alpha)$-grating. For our purposes, it is thus sufficient to tabulate the response of a vertical-preferring local population to gratings with different orientations, which we do below, beginning with how we model the dynamics of single LGN cells under

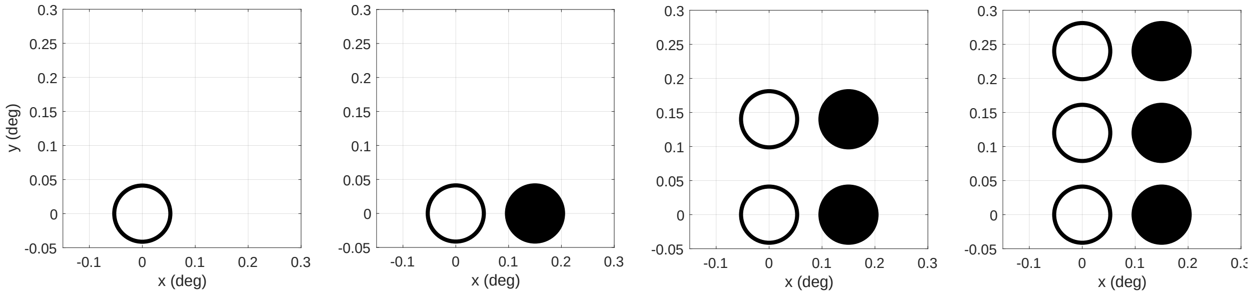


Figure S2: LGN configurations. White circles represent ON cells, and black circles represent OFF cells. The centers of the circles indicate the centers of LGN place fields, while the sizes of circles indicate the spatial standard deviations of place fields.

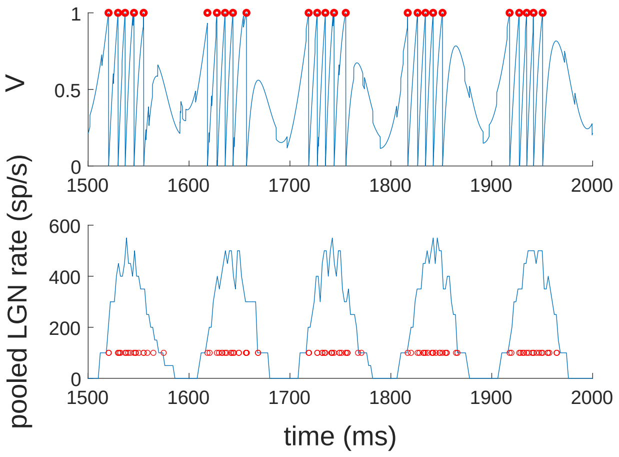


Figure S3: The responses of single LGN cells and pooled LGN responses when optimally driven. Top: the V trajectory of one ON LGN cell (blue) with spiking times (red circles). Bottom: the rates of the summed LGN spikes from all four LGN cells (blue) and the exact spiking pattern (red circles).

different visual stimuli, followed by the spatial configuration of LGN afferents and the resulting pooled input patterns to L4.

#### S2A Dynamics of single LGN cells

We adapt dynamic equations for single LGN cells from [3]. Our LGN cell model is another instance of the integrate-and-fire model:

$$\frac{dV}{dt} = -\frac{V}{\tau^{\text{leak}}} + I_{\pm}(t) + N(t), \tag{7}$$

where $V$ is membrane voltage; $\tau^{\text{leak}} = 10$ ms is the leakage timescale; $I_{\pm}(t)$ is the input due to visual stimuli presented in the receptive field of the LGN cell, with "+" and "−" indicating whether the LGN cell is "ON" or "OFF" (see below); and $N(t)$ is a Poisson pulse train modeling random noise:

$$N(t) = f \sum_i b^i \cdot \delta(t - t^i),$$

with $f = 0.075$, $b^i = \pm 1$ determined by independent tosses of a fair coin, and $\{t^i\}$ a Poisson process with rate $\lambda = 100$ Hz. When no visual stimuli are presented, $I_{\pm}(t) \equiv I_B = 100s^{-1}$, representing background input. (The unusual unit for currents is a consequence of our nondimensionalizing voltage but not time.) The firing threshold $V^{\text{th}}$ is set to 1.002 to ensure the background firing rate of a single LGN cell is approximately 20 Hz.

When a visual stimulus $L(\vec{x}, t)$ is presented to an LGN cell whose receptive field is centered at $\vec{x}_0$, the input current is given by

$$I_{\pm}(t) = [I_B \pm Q(t)]^+ \qquad \text{and} \tag{8}$$

$$Q(t) = Q_0 \int_0^t ds \cdot \left( \iint_{\vec{x} \in \mathbb{R}^2} K(s) A(\vec{x}_0 - \vec{x}) \cdot L(\vec{x}, t - s) \, d\vec{x} \right), \tag{9}$$

where $[x]^+ = \max(x, 0)$. Here, $Q(t)$ is the "visual gain" of the LGN cell from its receptive field due to the stimulus. In Eq. (9), $Q_0 = 0.58$ is a constant, and $K(t)$ and $A(\vec{x})$ are the temporal and spatial kernels of the LGN cell:

$$K(t) = \frac{t^6}{\tau_0^7} e^{-t/\tau_0} - \frac{t^6}{\tau_1^7} e^{-t/\tau_1}, \tag{10}$$

$$A(\vec{x}) = \frac{a}{\pi \sigma_a^2} e^{-|\vec{x}|^2/\sigma_a^2} - \frac{b}{\pi \sigma_b^2} e^{-|\vec{x}|^2/\sigma_b^2}. \tag{11}$$

The temporal and spatial kernels are identical for both ON and OFF cells. In the above, we take $(\tau_0, \tau_1) = (3.15, 6.17)$ ms, $(\sigma_a, \sigma_b) = (0.0894, 0.1259)^\circ$, and $(a, b) = (1.0, 0.74)$. These choices are based on [1]; the kernels are depicted in Fig. S4.

Finally, the visual stimulus $L(\vec{x}, t)$ represents the light intensity map for a drifting grating:

$$L(\vec{x}, t) = L_0 \cdot \left[ C \sin(-2\pi \vec{g} \cdot \vec{x} + 2\pi f t + \phi) + 1 \right], \tag{12}$$

where

– $C \geqslant 0$ is the contrast, with $C = 100\%$ being full contrast; in this paper, $C$ is usually set to a value between 0 and 64%;;

– $\vec{g} = (g_1, g_2)^T$ is a vector whose magnitude represents spatial frequency (in cycles/degree, always set to $2.5c/d$ in this paper) and whose direction is $(\cos\theta, \sin\theta)$, corresponding to a grating of orientation $\theta$ (with $\theta = 0$ being vertical);

– $f$ is the temporal frequency (in Hz), ranging from 2 to 24Hz in this study;

– $\phi$ is an overall phase, here set to 0; and

– $L_0 = 1$ is the mean light intensity, here set to 1.

The temporal kernel $K(t)$ effectively estimates the derivative of a spatial average of $L(\vec{x}, t)$, so that $Q(t)$ (and hence $I_+(t)$) is positive when the grating transitions from dark to bright within the LGN cell's receptive field; the opposite occurs for OFF cells. See [3] and references therein for more details.

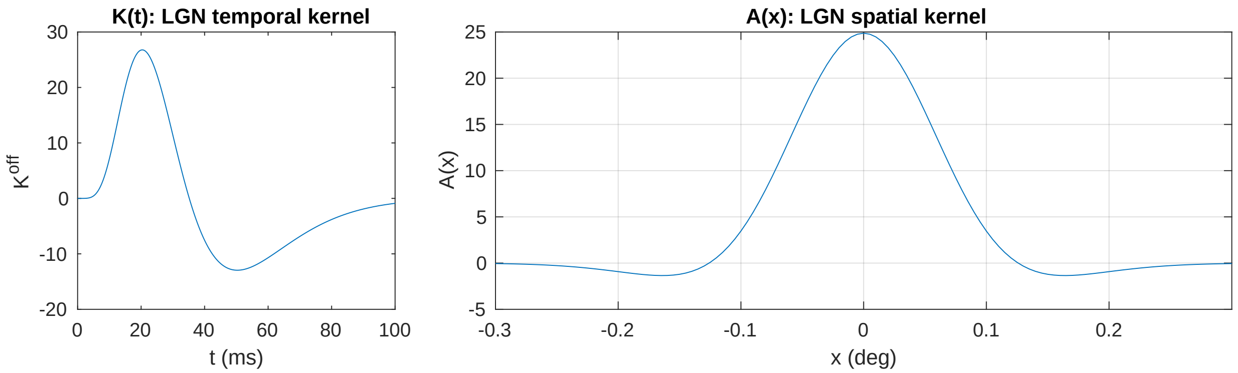


Figure S4: Kernels for LGN cells.

#### S2B Spatial configuration of LGN cells

Four spatial configurations of LGN cells are used to simulate LGN inputs to V1 cells: 1-cell, 2-cell, 4-cell, and 6-cell configurations. These configurations are designed to preferentially respond to vertical gratings (0° orientation).

– The 1-cell configuration centers the place field of an ON cell at $\vec{x}_{1,\text{on}} = (0, 0)^T$;

– The 2-cell configuration adds an OFF cell with its place field centered at $\vec{x}_{1,\text{off}} = (0.15, 0)^T$. Together, these form an ON-OFF pair;

– The 4-cell configuration introduces an additional ON-OFF pair at $\vec{x}_{2,\text{on}} = (0, 0.14)^T$ and $\vec{x}_{2,\text{off}} = (0.15, 0.14)^T$;

– The 6-cell configuration extends the 4-cell configuration with two additional ON-OFF pairs at $\vec{x}'_{2,\text{on}} = (0, 0.12)^T$, $\vec{x}'_{2,\text{off}} = (0.15, 0.12)^T$, $\vec{x}'_{3,\text{on}} = (0, 0.24)^T$, and $\vec{x}'_{3,\text{off}} = (0.15, 0.24)^T$.

The four LGN configurations are illustrated in Fig. S2.

##### S2C LGN inputs to L4 cells

We now describe how we generate LGN inputs to L4 cells. Recall the responses of $\alpha$-preferring orientation domains (ODs) for $\alpha = 45, 90, 135°$ can be equated to that of the $0°$-preferring OD to grating orientation $\theta - \alpha$; hence we focus on the vertical-preferring OD under drifting grating stimulation $L(\vec{x}, t)$. In our study, all visual stimuli are either background ($C = 0$ in Eq. (12)) or drifting gratings with orientations that are multiples of $7.5°$, i.e., $\vec{g} = g_0(\cos\theta, \sin\theta)^T$, where $g_0$ is the spatial frequency, and $\theta = n \times 7.5°$ for integer $n$.

For each LGN configuration, LGN cell dynamics are simulated for $T^{\text{LGN}} = 100$ s with a timestep of $\delta t = 0.1$ ms. For each $\theta$, four time series are obtained from the simulation: $\{t_i^1\}_\theta$, $\{t_i^2\}_\theta$, $\{t_i^4\}_\theta$, $\{t_i^6\}_\theta$, where the superscript denotes the number of LGN cells in the configuration. These time series are segmented into 100 sections of 1s each, which are used to construct LGN inputs for S and C cells.

**Simple cells.** Following [1–3], on average each S cell receives inputs from 4.5 LGN cells. Accordingly, we compute the firing rates of S cells by averaging those of S cells receiving LGN inputs from 4-cell and 6-cell configurations, i.e., for fixed L4E, L4I, and L6 inputs, we simulate two S-cells using Eq. (4), each for $T^{\text{L4}} = 20s$:

- one cell receives LGN input consisting of twenty randomly selected 1-s sections from $\{t_i^4\}_\theta$, as well as L4E, L4I, and L6 inputs as Poisson series with rates $(n^{SE}, n^{SI}, n^{S\text{L6}})$; and
- the other cell receives LGN input consisting of twenty randomly selected 1-s sections from $\{t_i^6\}_\theta$, as well as L4E, L4I, and L6 inputs as Poisson series with rates $(n^{CE}, n^{CI}, n^{C\text{L6}})$.

Let $f^{S,\text{4LGN}}$ and $f^{S,\text{6LGN}}$ denote the mean firing rates from the two simulations. The S firing rate is then defined to be $f^S = pf^{S,\text{4LGN}} + (1-p)f^{S,\text{4LGN}}$, where $p = 0.75$.

**Complex cells.** On average, one C cell receives inputs from 1.5 LGN cells [1–3]. Like S cells, the firing rates of C cells are averages of two C-cell simulations, with LGN inputs from 1- and 2-cell configurations, respectively. We use $f^C = qf^{C,\text{1LGN}} + (1-q)f^{C,\text{2LGN}}$ with $q = 0.5$.

**Inhibitory cells.** The firing rates of I cells are determined differently: following [1], we assume the LGN cells projecting to I cells are not configured to strongly prefer any particular orientation, so that LGN-to-I inputs are less regular and do not vary with $\theta$. To implement this feature, we construct LGN-to-I inputs in two steps: (i) we divide each 1s LGN spike train $\{t_i^4\}_{\theta'}$ into $\sim f$ equal segments, each lasting $\sim 1/f$ sec, where $f$ is the temporal frequency of drifting gratings; and (ii) for any grating orientation $\theta$, a 20s LGN input is formed by a random combination of sections from $\{t_i^4\}_{\theta'}$ where $\theta' \in \{45, 52.5, 60, \ldots, 90\}°$. The output firing rate of I cell $f^I$ is obtained from the 20s simulation receiving the constructed LGN input, as well as L4E, L4I, and L6 inputs as Poisson series with rates $(n^{IE}, n^{II}, n^{I\text{L6}})$.

#### S3 Coarse-grained model: implementation details

In this section, we describe how we compute the local response function, then use it to iteratively compute steady states for the CG model. Our focus here is to briefly summarize key points from [4] and highlight differences between this paper and our earlier work; interested readers are referred to [4] for further details.

##### S3A Constructing and evaluating local response function

The precomputation of local response functions, i.e., the construction of the "library" of local responses, is based on [4] with several modifications. Recall that type-$Q$ neurons ($Q \in \{S, C, I\}$) receive input from LGN, L4E, L4I, L6, and ambient sources. Here we fix the rate of ambient input. For a pixel $p$ lying in a vertical-preferring orientation domain, the firing rate $f_p^Q$ of type-$Q$ cells in $p$ in response to a grating of orientation $\theta$ is written

$$f_p^Q = \Phi_\theta^Q \left( n_p^{QE}, n_p^{QI}, n_p^{QL6} \right).$$

In the above, $\Phi_\theta^Q$ denotes the local response function type-$Q$ cells in $p$ to a $\theta$-grating, with all other stimulus parameters, e.g., contrast, light intensity, *etc* (see Sect. S2A), held fixed. The inputs of $\Phi_\theta(\cdot)$ are

- $n_p^{QE}$: combined spike rate from presynaptic L4 E-cells to each type-Q cell in pixel $p$;
- $n_p^{QI}$: combined spike rate from presynaptic L4 I-cells to each type-Q cell in pixel $p$; and
- $n_p^{QL6}$: combined spike rate from presynaptic L6 E-cells to each type-Q cell in pixel $p$.

As described in Sect. S2C, we precompute $\Phi_\theta^Q$ for selected angles $\theta$ for each type $Q$.

In [4], response functions were precomputed on overlapping 2D grids in the space of potential L4 excitatory (L4E) and inhibitory (L4I) input firing rates; on each 2D grid, inputs from LGN and L6 were fixed. In this study, due to the dependence of L6 input on L4 activity, we extend the precomputed library to 3D grids in the space of L4E, L4I, and L6 input rates. On each 3D grid, the LGN input is fixed and computed using simulated LGN cells (see Sect. S2); a range of possible LGN spike patterns are simulated prior to precomputing local response functions.

The 3D grids used in this paper are Cartesian products of the 2D grids used in [4] and a 1D grid of L6 firing rates. On the latter grid, L6 input is discretized into values ranging from 3Hz to 120Hz, in steps of 3Hz. For each L6 value, precomputation was carried out on the space of L4E and L4I inputs using the same strategy as described in [4].

Finally, the output $f_p^Q$ is determined by collecting the number of firing events from a 20-second simulation (excluding the first two seconds) of Eq. (4) using a timestep $\delta t = 0.1$ ms. Here, $\{t_k^{\text{lgn}}\}$ comes from the LGN input spiking pattern, whereas $\{t_k^{\text{amb}}\}$, $\{t_k^{\text{L6}}\}$, $\{t_k^{\text{L4E}}\}$, and $\{t_k^{\text{L4I}}\}$ are generated from independent Poisson processes with rates of $F^{Q\text{amb}}$, $n^{Q\text{L6}}$, $n^{QE}$, and $n^{QI}$, respectively.

**Evaluating precomputed local response functions.** Solving for steady state firing rates in the CG model will require repeatedly evaluating local response functions for different combinations of inputs. To do so, we interpolate $\Phi_\theta^Q$ from precomputed values of $\Phi_\theta^Q$ at neighboring grid points using MATLAB's `interp3` function with the "linear" option [4], which linearly interpolates between precomputed values along each coordinate.

##### S3B CG iterations

We now describe an iterative procedure for finding steady-state firing rates for the CG model. We denote the firing rates at step $n = 0, 1, 2, \cdots$ by $\left(\{f_p^{S,n}\}, \{f_p^{C,n}\}, \{f_p^{I,n}\}\right)$.

**Iteration scheme.** Following [4], in each iteration we replace the firing rates of each pixel with a mixture of its current firing rates and the pixel's local response to inputs from other pixels for a suitable stepsize $h \in (0, 1)$ tuned to balance convergence rate and stability [4]. We now describe how we compute the response.

**The "no-mixture" assumption.** For most L4 cells in one OD away from OD boundaries, their LGN inputs are essentially identical. The relatively small number of cells near OD borders may, however, have different response functions. Precomputing different local responses for cells away from and near borders is in principle possible, but more expensive, with questionable accuracy gains. Here we take first simplify the cortical model by imposing a "no-mixture" assumption that neglects the potentially different responses near borders, i.e., (i) each pixel is entirely contained within a single orientation domain; and (ii) LGN inputs are identical for all type-Q cells within each OD. While this assumption simplifies computation, LGN inputs do cross OD boundaries in reality. We then remove this "no-mixture" assumption and account for responses near boundaries by a suitable modification of the CG model. Note that LGN projections do not cross ODC borders; we will account for this below.

**L4 inputs.** As described in Sect. S1A, the combined spike rates of recurrent input to pixel $p$ ($n_p^{QR}$) sum the firing rates of E-cells around pixel $p$ through the L4 synaptic interaction kernel $C_{Q\leftarrow E,p\leftarrow p'}$:

$$n_p^{QE,n} = \sum_{p'\in\mathscr{P}} C_{Q\leftarrow E;p\leftarrow p'}\, f_{p'}^{E,n}. \tag{13}$$

Inhibitory inputs are computed differently: as discussed in Sect. S1A, inhibitory inputs are assumed to be depressed at high firing rates. We implement this as follows: for each pixel, the inhibitory firing rate $f_p^{I,n}$ is replaced by $\hat{f}_p^{I,n} = f_p^{I,n}\cdot s(f_p^{I,n})$, where $s(\cdot)$ is a depression factor ranging between 0 and 1. (See Fig. S1.) The combined I-spike rate to pixel $p$ is thus

$$n_p^{QI,n} = \sum_{p'\in\mathscr{P}} C_{Q\leftarrow I;p\leftarrow p'}\, \hat{f}_{p'}^{I,n}. \tag{14}$$

**L6 inputs.** Unlike [4], where the L6 input was static, here we dynamically update L6 input based on local L4E firing rates, using a kernel $k$; see Eq. (2) in Sect. S1A. The combined L6E-spike rate to pixel $p$ is

$$n_p^{QL6,n} = m^{QL6}\cdot R\Big(\sum_{p'\in\mathscr{P}} k_{p\leftarrow p'}\, f_{p'}^{E,n}\Big), \tag{15}$$

where $m^{QL6}$ is the average number of presynaptic L6 cells to a type-$Q$ L4 cell (see Sect. S1C), and $R(\cdot)$ is the L6 response function shown in Fig. S1.

**Averaging outputs for different LGN inputs.** We now relax the no-mixture assumption made earlier. To account for responses near borders, we follow [4] and average the *output firing rates* of nearby pixels, rather than compute separate response functions for cells near borders. To aid in assigning weights to nearby pixels, we first ascribe the probability that an LGN cell at $\mathbf{x}'$ is connected to an L4 cell at $\mathbf{x}$; this probability is not used elsewhere. How we do this depends on whether the model in the single and two-eye models:

- **Preliminary (single-eye) model:** All LGN inputs are assumed to originate from one eye. The probability that an LGN cell at $\mathbf{x}'$ projects to an L4 cell at $\mathbf{x}$ is modeled by a truncated Gaussian $P^{\mathrm{LGN}}(\mathbf{x}'-\mathbf{x})$ with spatial standard deviation $\sigma = 100\,\mu$m and truncation radius $r = 100\,\mu$m. $P^{\mathrm{LGN}}$ is not affected by the boundaries between HCs. For local population $p$, the weight of LGN input from $\alpha$-preferring OD is

$$w_p^{\alpha} = \int_{\mathbf{x}\in\text{pixel } p} d\mathbf{x} \int_{\mathbf{x}'\in OD_\alpha} P^{\mathrm{LGN}}(\mathbf{x}'-\mathbf{x})\, d\mathbf{x}'. \tag{16}$$

The firing rate maps $\{f_p^{S,n}\}$, $\{f_p^{C,n}\}$, and $\{f_p^{I,n}\}$ are then updated according to

$$f_p^{Q,n+1} = (1-h) f_p^{Q,n} + h \sum_\alpha w_p^\alpha \Phi_{\theta-\alpha}^Q \left(n_p^{QE,n}, n_p^{QI,n}, n_p^{\mathrm{QL6,n}}\right) \tag{17}$$

in each step of the iteration.

– **Two-eye model:** It is well known that LGN cells from one eye do not project to ODCs of the other eye. To model this (and ensure L4 cells near ODC boundaries do not have artificially low numbers of LGN afferents in our model), we modify $P^{\mathrm{LGN}}$ by sending LGN projections that would have crossed ODC boundaries back to the home ODC. Note this "trick" is only to maintain LGN input to L4 cells; in reality LGN-to-L4 projections are not folded back. The adjusted probabilities are

$$\hat{P}^{\mathrm{LGN}}(\mathbf{x}, \mathbf{x}') = \begin{cases} P^{\mathrm{LGN}}(\mathbf{x}' - \mathbf{x}) + P^{\mathrm{LGN}}(\mathbf{x}'' - \mathbf{x}) & \text{if } \mathbf{x}' \text{ is in the same ODC as } \mathbf{x}, \\ 0 & \text{if } \mathbf{x}' \text{ is in a different ODC,} \end{cases} \tag{18}$$

where $\mathbf{x}''$ is the reflection of $\mathbf{x}'$ along the closer of the two boundaries for its ODC. (In our model, no L4 cell is within 100$\mu$m of both ODC boundaries, so that this rule is sufficient to maintain the number of LGN afferents.) Weights $w_p^\alpha$ are then computed using $\hat{P}^{\mathrm{LGN}}$, and the firing rate map is updated according to

$$f_p^{Q,n+1} = (1-h) f_p^{Q,n} + h \sum_\alpha \hat{w}_p^\alpha \Phi_{\theta-\alpha}^Q \left(n_p^{QE,n}, n_p^{QI,n}, n_p^{\mathrm{QL6,n}}\right)$$
$$\hat{w}_p^\alpha = \int_{\mathbf{x} \in \text{pixel } p} d\mathbf{x} \int_{\mathbf{x}' \in OD_\alpha} \hat{P}^{\mathrm{LGN}}(\mathbf{x}, \mathbf{x}')\, d\mathbf{x}'.$$

**Additional details.** Eachc simulation of the basic CG model and the two-eye model consists of 50 iterations. This number is sufficient for both models to converge to a stable fixed point. The firing rate maps of the final iteration typically exhibit an "HC"-norm difference of $o(0.1\text{Hz})$ from the fixed point (see the definition of HC-norm in [4]). This choice is also influenced by computational resource constraints. For reference, for our simulations — which were implemented using MATLAB (2022a) from MathWorks running on a 16-core machine (with Advanced Micro Devices Ryzen 9 CPUs running Ubuntu 20.04 LTS) — a single 50-iteration simulation of the two-eye model requires approximately 18 seconds. To generate the results presented in the main text, at least 250 simulations of the basic CG model and 675 simulations of the two-eye model were conducted.

### S4 Supplemental Analysis

#### S4A Analysis of cross-ODC cortical currents in the CG model

In our analysis of the binocular model, we asserted that under monocular stimulation, L4 currents received by local populations via cross-ODC connections are net-negative. This played a key role in our discussion of Findings 1-3. Here we present concrete evidence supporting this assertion.

**Decomposition of currents to L4E neurons.** First, as the CG model has no explicit representation of synaptic currents, we need to estimate the cortical currents received by excitatory subpopulations within each local population. To do so, we begin by computing the rates of incoming L4E and L4I inputs from steady-state firing rate maps $\left(\{f_p^{S,*}\}, \{f_p^{C,*}\}, \{f_p^{I,*}\}\right)$ of the CG model. Then we simulate LIF neurons (Eq. (4)) and compute the mean current from type-$R$ cell to type-$Q$ cell by

$$\frac{1}{T}\int_{\mathbb{T}} g_R^Q(t)(V(t)-V^R)\,\mathrm{d}t\ , \tag{19}$$

where $Q \in \{S, C\}$, $R \in \{E, I\}$, and $\mathbb{T}$ is a time window of duration $T$ excluding refractory periods. Here we use $T = 20$ seconds. For a local population $p$, we then estimate the total L4 input currents to E-subpopulation as weighted averages of currents to S and C subpopulations, i.e.,

$$\text{total L4E current} = a^S \cdot \text{L4E current to an S-cell} + a^C \cdot \text{L4E current to a C-cell},$$
$$\text{total L4I current} = a^S \cdot \text{L4I current to an S-cell} + a^C \cdot \text{L4I current to a C-cell},$$

where $a^S$ and $a^C$ are the fractions of S and C cells within the E-subpopulation.

Fig. S5 shows the current decomposition for a CG model consisting of $2 \times 2$ hypercolumns, with the upper and lower HCs belonging to distinct ODCs. The top panels (A-C) show the response to a vertical grating presented to both eyes; the bottom panels show the response to monocular stimulation by a 45° grating. In both settings we see that L4 currents (right column, i.e., panels C and F) are net negative.

**Current analysis for Finding 1.** We had concluded that $\mathbf{P}(Abs)$ (the probability that potential cross-ODC connections are deleted) is small to negligible, on the basis that unrealistically high firing rates are seen near ODC boundaries with appreciable values of $\mathbf{P}(Abs)$. This elevated activity was attributed to net negative cross-ODC currents from within L4. To see this, first consider Fig. 4C in the main text: had all connections been reflected (top right panel), L4 inputs received by the home column of a local population during monocular stimulation would have been identical to that during binocular stimulation. But when $\mathbf{P}(Abs) = 1$ (top left panel), cross-ODC connections are deleted rather than reflected, causing local populations adjacent to the border to miss nearly half of their input currents (both E and I) from L4. Fig. S5C confirms that these "missed" currents are indeed net-negative, leading to elevated firing rates. There is also current loss in the second row from the border, though by a smaller amount.

**Current analysis for Finding 3.** Fig. S5E shows the input currents from within L4 when the top ODC alone is stimulated. For the emergence of binocular strips near ODC borders (Finding 3 in the main text), we are interested in other-eye stimulation, which in the context of Fig. S5E means the response of local populations immediately below the ODC border. Observe (i) net L4 currents in panel F are net-negative, and (ii) in panel D, the elevated firing rates induced by strong positive currents from L6, which is in turn due to the high firing just across the border in the stimulated ODC. (Recall that L6 activity is locally indexed to that of E subpopulations in L4, and their projections have a range of $3 \times 3$ pixels, thus reaching 1 pixel into the unstimulated ODC.)

#### S4B Two-parameter family of CG models

Fig. S6 shows firing rate maps for 45°-gratings, under different choices of $\mathbf{P}(Crs)$, $\mathbf{P}(Rfl)$, and $\mathbf{P}(Abs)$. This complements Fig. 4C in the main text (which shows the response to a vertical grating). As in Fig. 4C in the main text, $\mathbf{P}(Abs) > 0$ leads to notable artifacts near ODC boundaries. Note also the abnormally high E-firing rate of pixels of the unstimulated ODC close to OD boundaries (in light blue). These observations further support our conclusion that $\mathbf{P}(Abs) \approx 0$ and $\mathbf{P}(Crs) \in [0.25, 0.50]$.

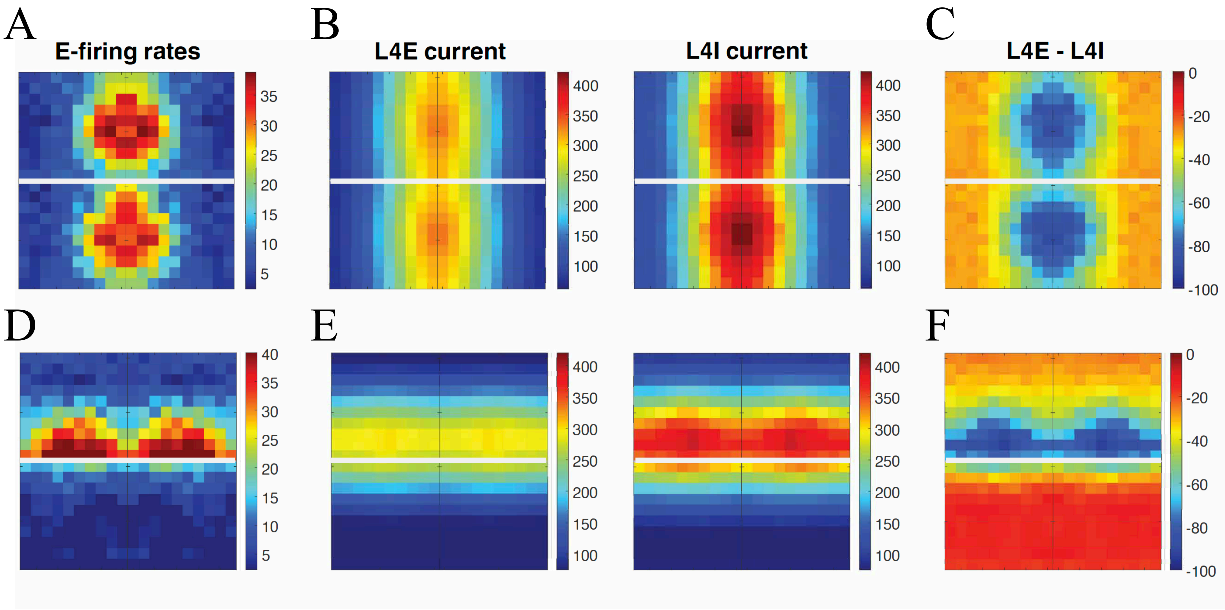


Figure S5: Current decomposition. Here we drive a $2 \times 2$-HC CG model with different gratings, at full contrast with a temporal frequency of 10 Hz. The upper and lower HCs belong to the ODCs of the left and right eyes, respectively. Top row (panels A-C): we present a vertical grating to both eyes. Panel (A) shows the resulting E firing rates. Panel B: E and I currents received by E populations from within L4. Panel C: net L4 currents. Observe L4 currents are net-negative throughout, and is more negative in areas of higher activity. Bottom row (panels D-F): a 45° grating is presented to the eye corresponding to the top ODC. Observe the elevated firing rates of pixels immediately below the ODC border, due to strong L6 feedback. Again, L4 currents (panel F) are seen to be net-negative.

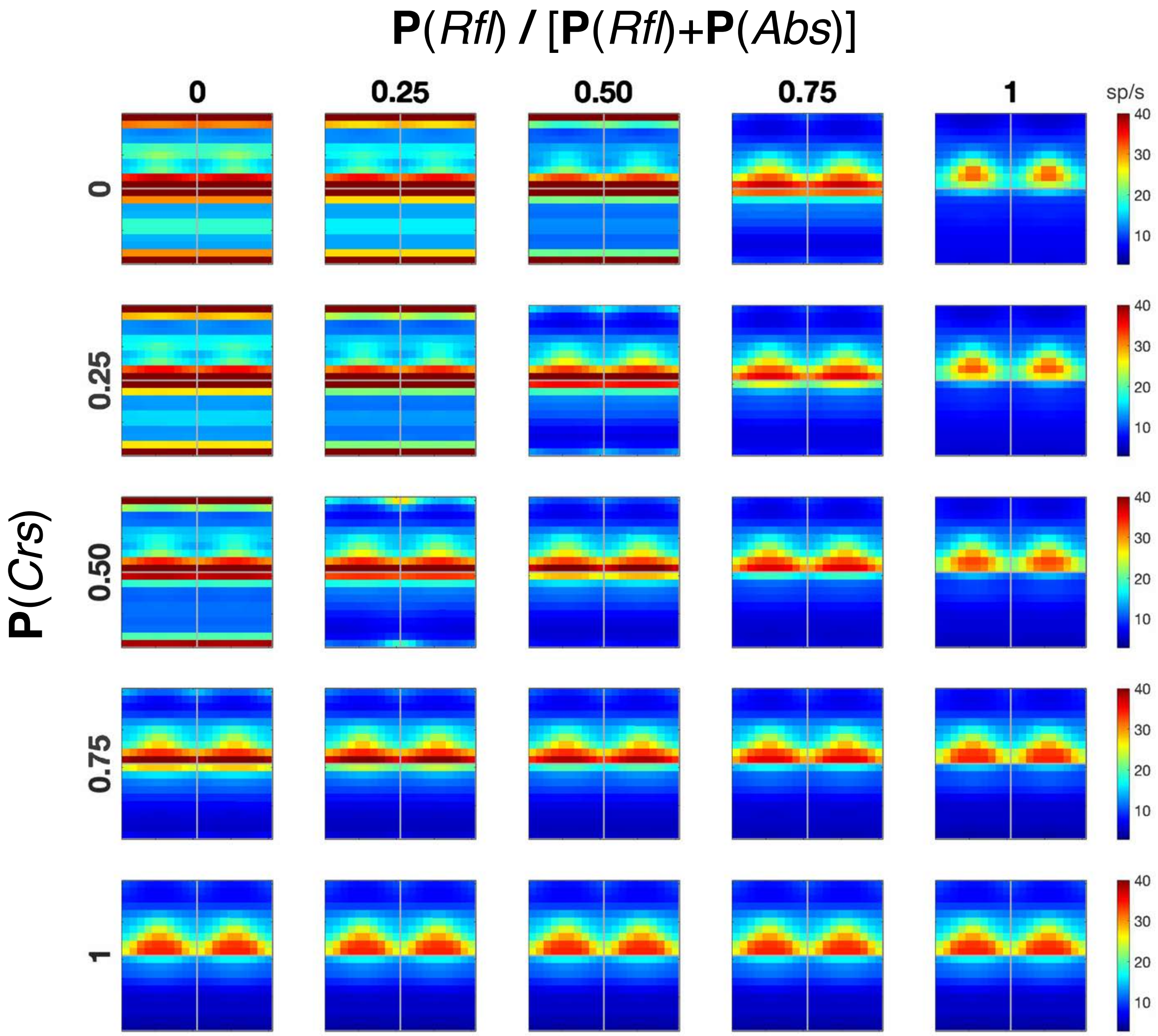


Figure S6: Firing rate maps for 45°-gratings under different choices of **P**(*Crs*), **P**(*Rfl*), and **P**(*Abs*). This complements Fig. 4C in the main text.

#### S5 Binocularity of L4 and L6

In Figs. 6–8 of the main text, we introduced various measures designed to quantify the degree to which cortical cells respond to binocular stimulus. Here we define these measures precisely and provide some additional information.

##### S5A Binocularity index of L4

The binocularity index (see Fig. 6) and binocular modulation (BM; Fig. 7) of a pixel $p$ depend on three tuning curves as functions of grating orientation $\theta$: $b_p(\theta)$, the E-firing rate of pixel $p$ when both eyes are stimulated; $m_p(\theta)$, the E-firing rate when only the dominant eye of pixel $p$ is stimulated; and $o_p(\theta)$, the E-firing rate when only the other eye is stimulated. We fix $\mathbf{P}(Abs) = 0$ and vay $\mathbf{P}(Crs)$, and compute L6 as described in the text (i.e., proportional to L4). The stimulus has temporal frequency 10 Hz, spatial frequency 2.5 c/d, and full contrast (i.e. 67%).

Let $f_{bg}$ be the background E-firing rate and $[r] = r - f_{bg}$, i.e., firing rate (in Hz) above background. To compute the binocularity index (BI), we first maximize the response $m_p(\theta)$ of pixel $p$ to monocular stimulation of its dominant eye over grating angle $\theta$; let $\theta_p^m = \arg\max_\theta m_p(\theta)$ be the grating angle that maximizes $m_p(\theta)$ and $\bar{m}_p = [m_p(\theta_p^m)]$, i.e., $p$'s maximum firing rate above background under monocular stimulation. Then we define

$$BI_p = \frac{\bar{o}_p}{\bar{o}_p + \bar{m}_p} \ , \quad \bar{o}_p = [o_p(\theta_p^m)] \ . \tag{20}$$

The BI is always nonnegative, and a larger $BI_p$ indicates $p$ responds more when the other eye is stimulated. Note that $1 - 2BI_p$ is the conventional binocularity measure used by Hubel and Wiesel and other authors [14–16].

To compute the binocular modulation BM of a pixel $p$, we now maximize the binocular response $b_p(\theta)$ over $\theta$, with $\theta_p^b = \arg\max_\theta b_p(\theta)$ the grating angle that maximizes $b_p(\theta)$ and $\bar{b}_p = [b_p(\theta_p^b)]$. Then

$$BM_p = \frac{\bar{b}_p - \bar{m}_p}{\bar{b}_p + \bar{m}_p} \ . \tag{21}$$

This definition is identical to that used in other recent studies of binocular modulation [15, 16]. A positive BM means the pixel is more responsive to binocular stimulation than to monocular dominant-eye stimulation.

##### S5B L6 binocularity

In Fig. 8, we studied the degree to which L6-to-L4 feedback reflects information from both eyes. To test this, we first obtain the two steady-state L4E rate maps, $\{f_p^{E,b}\}$ and $\{f_p^{E,m}\}$, with $\{f_p^{E,b}\}$ the response to presenting the same grating to both eyes, and $\{f_p^{E,m}\}$ the response to stimulating only one eye. The corresponding binocular and monocular L6 inputs to $p$ are computed according to Eq. (15), i.e.,

$$b_6(p) = m^{QL6} \cdot R\Big(\sum_{p' \in \mathscr{P}} k_{p \leftarrow p'} f_p^{E,b}\Big) \qquad \text{and} \qquad m_6(p) = m^{QL6} \cdot R\Big(\sum_{p' \in \mathscr{P}} k_{p \leftarrow p'} f_p^{E,m}\Big) \ ,$$

where $R$ is the L6 response function (see Sect. S3B and Fig. S1). Setting L6 binocularity to $a \in [0,1]$ means that, in every iteration, the combined L6E-spike rate of pixel $p$, i.e., $n_p^{QL6}$ in Eq. (15), is replaced by the weighted average

$$\hat{n}_p^{QL6} = (1-a)\cdot m_6(p) + a\cdot b_6(p) \tag{22}$$

on every iteration. Note that unlike the algorithm described in Sect. S3B, in which L6 feedback is a linear function of L4 activity and thus updated whenever L4 rates are updated, here the quantities $m_6(p)$ and $b_6(p)$ are precomputed and held fixed during iteration.

##### S5C L6 binocularity at higher crossing probability

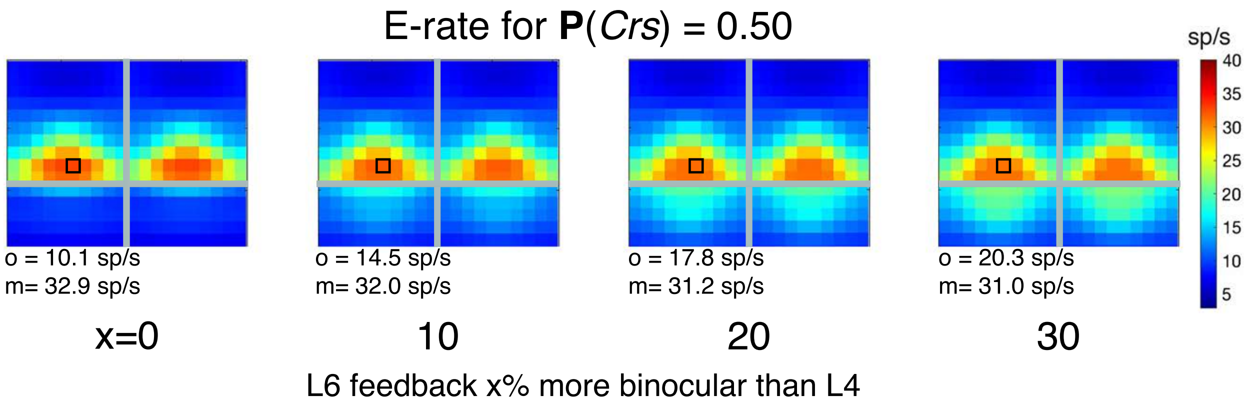


Figure S7: L6 binocularity for $P(Crs) = 0.5$. This complements Fig. 8 in the main text.

In Fig. 8 in the main text, we showed L6 binocularity with $\mathbf{P}(Crs) = 0.25$) and argued that this constrains the binocularity of L6 feedback to $< 10\%$ of L4. Fig. S7 shows L6 binocularity for $\mathbf{P}(Crs) = 0.5$. The degree of L6 binocularity in Fig. S7 is generally comparable to that of Fig. 8, and generally supports the same conclusion.
